# PCR-based detection of *Helicobacter* spp. in mice from different animal houses in Rio de Janeiro, Brazil

**DOI:** 10.1101/727420

**Authors:** Gabriel E. Matos-Rodrigues, Carolinne C. Masseron, Fabio J. Moreira da Silva, Marcel Flajblat, Lilian O. Moreira, Rodrigo A. P. Martins

**Affiliations:** Programa de Biologia Celular e do Desenvolvimento, Instituto de Ciências Biomédicas, Universidade Federal do Rio de Janeiro, Rio de Janeiro, RJ, Brazil; Faculdade de Farmácia, Universidade Federal do Rio de Janeiro, Rio de Janeiro, RJ, Brazil; Instituto de Biofísica Carlos Chagas Filho, Universidade Federal do Rio de Janeiro, Rio de Janeiro, RJ, Brazil

**Keywords:** Diagnostic, screening, laboratory mice, pathogen-free, infection

## Abstract

Pathogenic microbial detection and control in breeding and experimental laboratory animal facilities is essential to guarantee animal welfare, data validity and reproducibility. *Helicobacter spp.* is known to severely affect mice health, mainly in immunocompromised strains, what may affect experimental outcomes. This study aimed to screen for *Helicobacter spp.* in mice from four different animal houses in Rio de Janeiro, Brazil using a PCR for 16S ribosomal RNA. A pair of primers was designed to specifically identify *Helicobacter* species that commonly infect laboratory mice. Following PCR reaction, the expected 375 base pairs (bp) amplification product was purified, sequenced and showed a 95% similarity when compared to deposited sequences of *Helicobacter hepaticus* and *Helicobacter bilis*. Then, the presence of *Helicobacter spp.* in both feces and intestines samples was analyzed. *Helicobacter spp* DNA was detected in 59.6% of the fecal and 70.17% of the intestine samples. Although *Helicobacter spp* screening is recommended by institutional animal health monitoring programs worldwide it is still not mandatory by Brazilian animal welfare regulation. Our study, the first to monitor *Helicobacter* species in laboratory mice in Brazil, demonstrates the possibility of using a low cost, rapid molecular diagnostic test to screen *Helicobacter* and highlights the importance of regular microbiological verification of mice used for research in Brazilian animal houses.

## Introduction

The use of rodents as experimental models in basic and pre-clinical research has been essential for scientific progress. Reproducible research requires the use of laboratory animals free of diseases and other conditions that could interfere with experimental outcome. Infections that naturally occur in mice, even when subclinical, may influence animal’s physiology, immunity and behavior. For this reason, even in the absence of clinical signs, experimental rodents may become inadequate for research due to the presence of microorganisms, like bacteria, viruses or protozoa (Baker et al. 1998; Nicklas et al. 1999; Whary and Fox 2006; Besselsen et al. 2008, Pritchett-Corning et al. 2009). Moreover, immunosuppressed mice are prone to exhibit clinical manifestations from natural infections (Whary and Fox, 2006; Chichlowski and Hale, 2009). Therefore, the establishment of animal health status monitoring routines and quality control of laboratory mice maintained in facilities is indispensable (FELASA, 2014).

*Helicobacter* bacteria are among the microorganisms that infect laboratory rodents (Pritchett-Corning et al. 2009). *Helicobacter* spp. colonizes primarily rodents’ cecum and colon, but also stomach, gallbladder and liver. These microorganisms are shaded in feces leading to horizontal transmission through fecal-oral contact (Whary and Fox, 2006). Previous studies reported that, in mice, *Helicobacter* spp. infection is associated gastrointestinal and inflammatory bowel diseases as well as breast, liver, gastric and colon cancers (Chichlowski and Hale 2009) and that it can also affect mice reproduction (Sharp et al. 2008; Chichlowski and Hale 2009). In addition, immunodeficient animals are more susceptible to *Helicobacter* infection. In these mice, infection by *H. hepaticus, H. bilis, H. muridarum H. rappini* may lead to intestine manifestations, such as thyphlocolitis, hepatitis, gastritis and cancer (Whary et al. 2006; Mähler and Nicklas, 2012).

*Helicobacter spp.* are highly prevalent bacteria in animal facilities worldwide (Taylor et al. 2007; Wasimuddin et al. 2012) and were previously detected in wild rodents in Brazil (Comunian et al., 2006). To our knowledge, the prevalence of *Helicobacter* infection in laboratory rodents in Brazil has never been studied. The present study aimed to detect *Helicobacter spp.* bacteria in rodents of animal houses localized at Universidade Federal do Rio de Janeiro, Brazil using a PCR-based diagnostic test.

## Materials and methods

### Mice

The mouse strains were obtained from four different animal houses (AH-A–AH-D) localized at Universidade Federal do Rio de Janeiro, Brazil. Mice of mixed genetic background (n= 57) between 15 and 22 weeks of age (mean age, 19 weeks) were randomly selected. All animals were housed conventionally, fed a standard diet and water ad libitum, and euthanized by exposure to CO_2_ prior to intestine (cecum and/or colon) collection. Recovered feces (1 fecal pellet per animal) and intestine sections were frozen in liquid nitrogen and maintained at −80°C. All mice were treated according to the protocol (No. 092/15) approved by the Animal Ethics Committee of UFRJ.

### DNA extraction

DNA extraction of fecal samples (about 1 cm pellet, ~50-80 mg) (Beckwith et al., 1997) or lower bowel (about 1cm of cecum or colon) was performed using two different methods. In method one, samples were incubated in 100μL of 25 mM NaOH (Isofar, 1326) for 1h at 98°C. Then, 400μL of 10 mM pH 7.4 Tris buffer (Sigma, T1503) was added and the samples were stored at −20°C. In method two, samples were incubated in 200μL proteinase K 1mg/mL (Sigma, P2308) for 16h at 50°C and then mixed with 160μL of saturated NaCl (Isofar, 310) following centrifugation at 19.000g for 15min. Then 300μL of the supernatant were collected, mixed with 1mL of cold ethanol (Merck, 100983), and centrifuged at 19.000g for 15min. DNA pellets were ressuspended in 200μL of TE buffer (Tris 10mM-EDTA 1mM, pH 7.6) and stored at −20°C.

### Polymerase chain reaction

Previously described primer sequences that recognize a conserved region of the 16S ribosomal RNA gene (16S rRNA) and specifically detect four *Helicobacter* species (*H. hepaticus, H. bilis, H. muridarum and H. rappini*) were used (Proietti et al. 2009; Bury-Moné et al. 2003; Beckwith 1997; Riley et al. 1996). The reaction mixture (final volume 20μL) contained 0.5 μM of each primer (H276f: 5’-CTATGACGGGTATCCGGC-3’ and H676r: 5’-ATTCCACCTACCTCTCCCA-3’), 5x Green GoTaq® Flexi Buffer (Promega M891A), 1.5 mM MgCl_2_ (Promega, A351H), 10mM DNTP (Fermentas, R0199) and 0.025 U of Hotstart Taq polymerase (Promega) and different amounts of DNA (from 0.01ng to 100ng). The amplification conditions used were 94°C for 5’ followed by 35 cycles of 2’ at 94°C, 2’ at 53°C and 30” at 72°C for 5’ at 72°C. The PCR product was analyzed by electrophoresis in 1% agarose gel (Sigma, A9539).

### DNA sequencing

The 375 bp PCR amplicon was purified (GE Healthcare Illustra GFX PCR DNA and Gel Band Purification kit (GE, 28903470) according to the manufacturer’s instructions. A 7μL of the reaction mixture, containing 50ng of DNA template and 3.2 pmol of primer (5’-CTATGACGGGTATCCGGC-3’) was submitted to an ABI 3130xl automated sequencer. Obtained sequence was compared to *H. hepaticus, H. bilis, H. muridarum, Mus musculus* and *Escherichia coli* genomes using the Blast platform (https://blast.ncbi.nlm.nih.gov).

## Results

### PCR assay and amplicon sequencing

To investigate the presence of *Helicobacter* spp, we performed PCR assays using primers previously designed to recognize a conserved region of the 16S rRNA gene, specific for the *Helicobacter* genus (Riley et al. 1996; Proietti et al. 2009; Bury-Moné et al. 2003; Beckwith, et al., 1997). Primer-BLAST tool was used to confirm primers annealing against three different genomes: (i) *H. hepaticus* (GI: 32263428), (Riley et al., 1996), (ii) *Mus musculus* (GI: 372099100), to search to nonspecific amplifiable targets and (iii) to *E. coli* (GI: 26111730), considering the possibly of non-specific annealing with DNA from a bacteria commonly found in feces. Primer-BLAST search confirmed that the selected primers specifically recognize a conserved region of the *H. hepaticus* genome, indicating that it would also anneal to the DNA of other *Helicobacter* species.

Since feces may contain substances that can inhibit the PCR and lead to false negative results (Monteiro et al. 1997), we first tested NaOH extracted DNA from the feces of a mouse naturally infected with *Helicobacter spp* (kindly provided by Dr. Rovilson Gilioli, CEMIB, Unicamp). A serial dilution (0.1, 1, 10, and 100ng) of this *Helicobacter spp.*-containing DNA was performed and, as expected from the bioinformatics analysis, only the expected amplicon of 375 bp was detected in the three higher DNA concentrations tested (Figure 1). The successful amplification using 1 or 10 ng of DNA indicated a reasonable sensitivity of this reaction. To evaluate primers specificity to *Helicobacter* species, we isolated *E. coli* bacteria and extracted its DNA using two different methods (NaOH or proteinase K). Following PCR, no amplification was observed with the input of 1, 10 or 100 ng of *E. coli* DNA (Figure 2).

**Figure 1:**
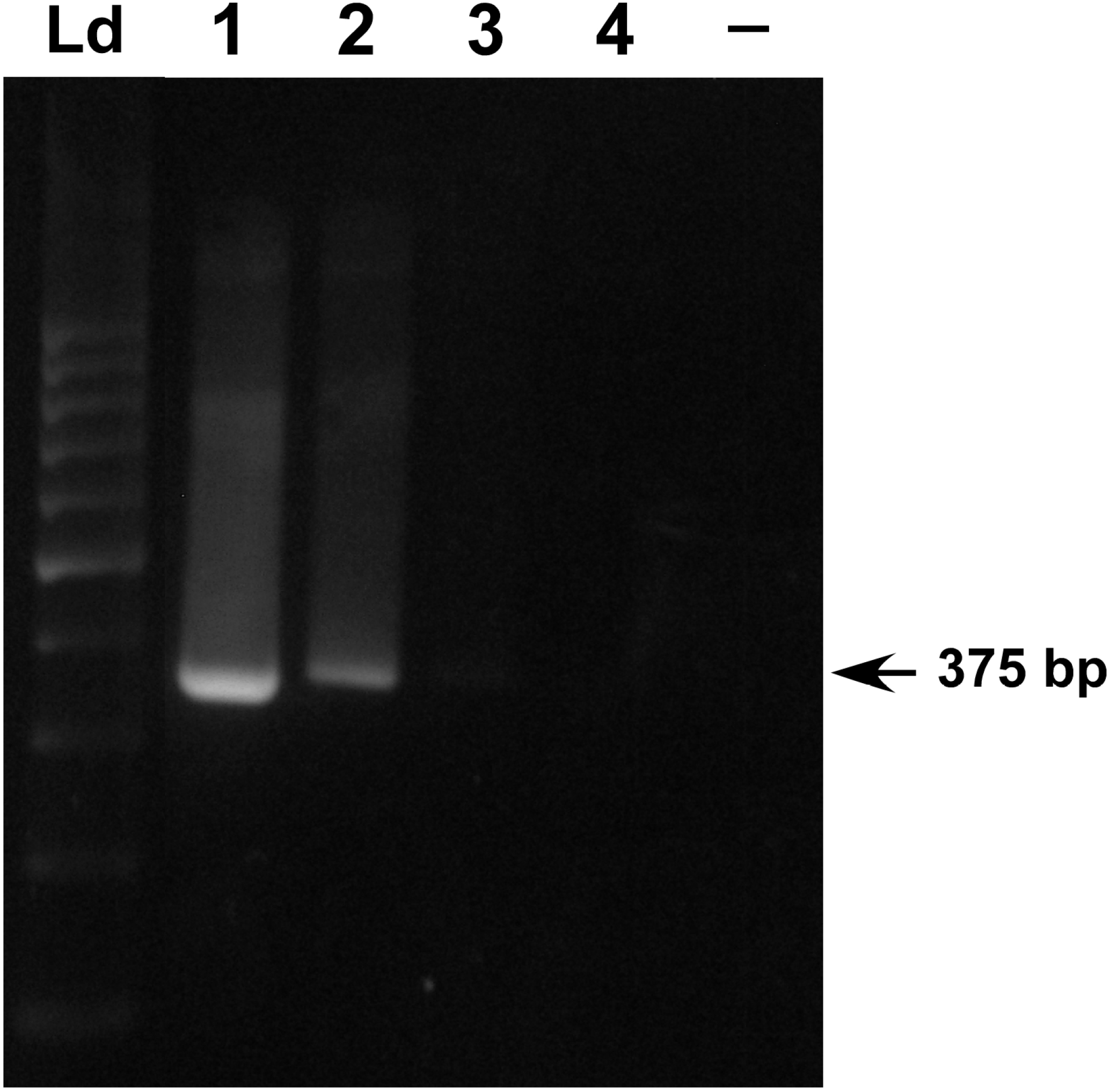
Sensitivity of the PCR-based assay for *Helicobacter spp.* detection. DNA extracted from feces of a mouse that was previously diagnosed with Helicobacter infection was used as the input of the PCR. Labels: Ld = 100 bp DNA ladder; 1 = 100 ng; 2 = 10 ng; 3 = 1 ng and 4 = 0.1 ng and (−) = no DNA; 1% agarose gel.

**Figure 2:**
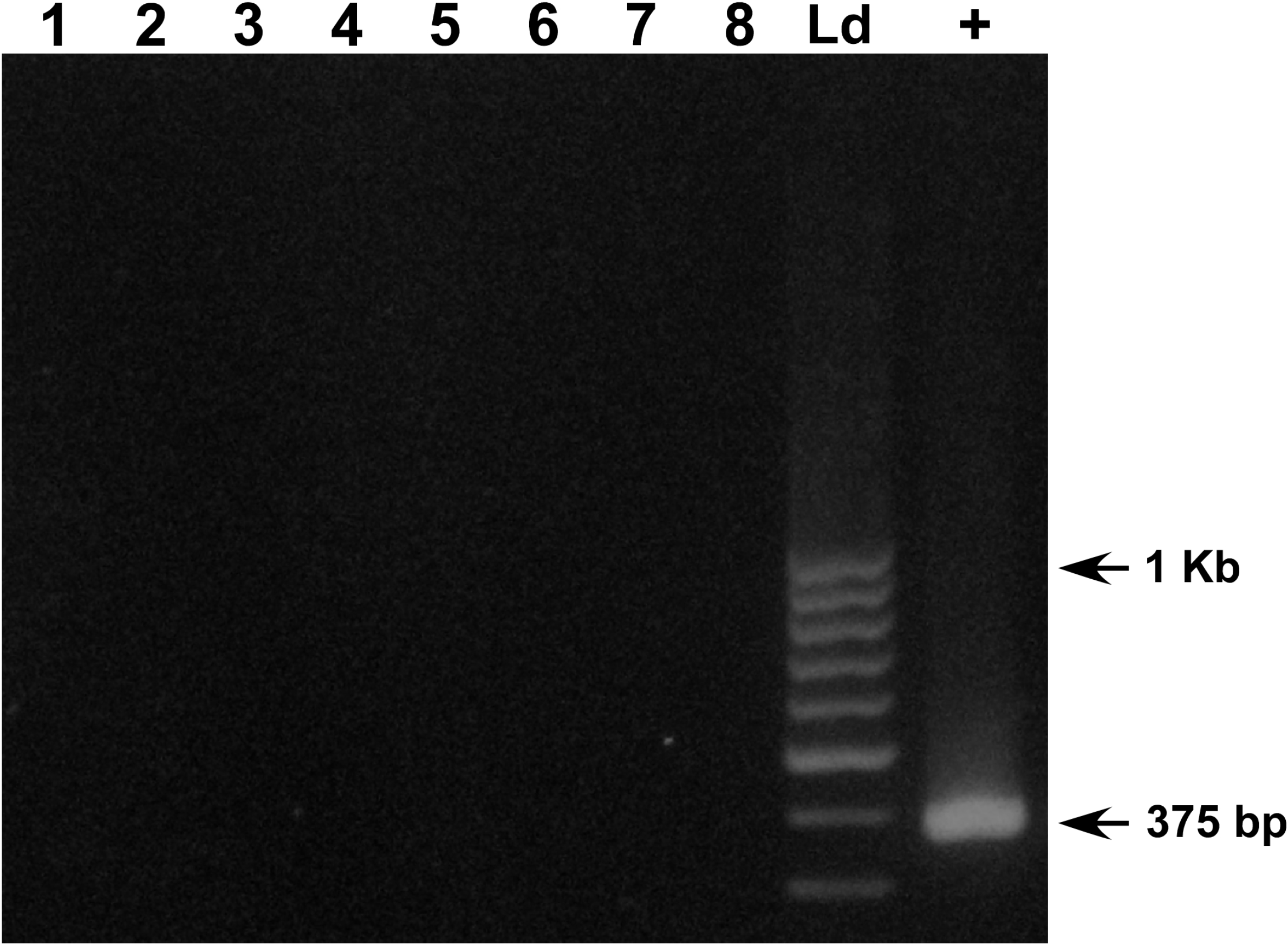
Specificity of the PCR-based assay for *Helicobacter spp.* detection. DNA extracted from *E. coli* bacteria (1-8) and DNA extracted from feces of a mouse that was previously diagnosed with Helicobacter infection (+) were used as the input of the PCR. Labels: Ld = 100 bp DNA ladder; 1-4: respectively, 100, 10, 1 or 0.1 ng of *E. coli* extracted with NaOH protocol; 5-8: respectively, 100, 10 1 or 0.1 ng of *E. coli* extracted with proteinase K protocol; (+) 100 ng of *Helicobacter spp.* DNA positive control; 1% agarose gel.

To determine whether the obtained PCR 375bp amplicon would correlate to the region of the *Helicobacter* genome, two samples were amplified, the PCR products were purified, sequenced and compared to the 16S rRNA gene sequences of *H. hepaticus* (GI:32265499), *H. bilis* (GI:806984271) and *H. muridarum* (GI:219846348) using BLAST platform. A ~95% similarity between the sequence of our amplicons and those deposited in NCBI database further validated the specificity of the PCR assay.

### PCR-based detection of *Helicobacter spp.* in mice

In order to evaluate the presence of *Helicobacter spp.* in mouse samples, first we collected feces or intestine DNA samples from 57 randomly selected mice from four different animal houses. Then, we performed PCR analysis to analyze the presence of *Helicobacter* spp DNA in these samples (Figure 3). We found that 59.6% of the feces samples were positive, while 70.17% (40 of 57) of the intestine samples contained detectable *Helicobacter* spp DNA (Table 1). It is important to point that not all animals were tested for both feces and intestines due to sampling limitations. According to the PCR-based diagnostic tests of the feces, the occurrence of *Helicobacter spp.* infection in the different animal houses (AH) ranged from 84% (AH-C) to 14% (AH-D). The analysis of intestine samples revealed a proportion of *Helicobacter spp.*-infected mice ranging from 90% (AH-B) to 55% (AH-D).

**Figure 3:**
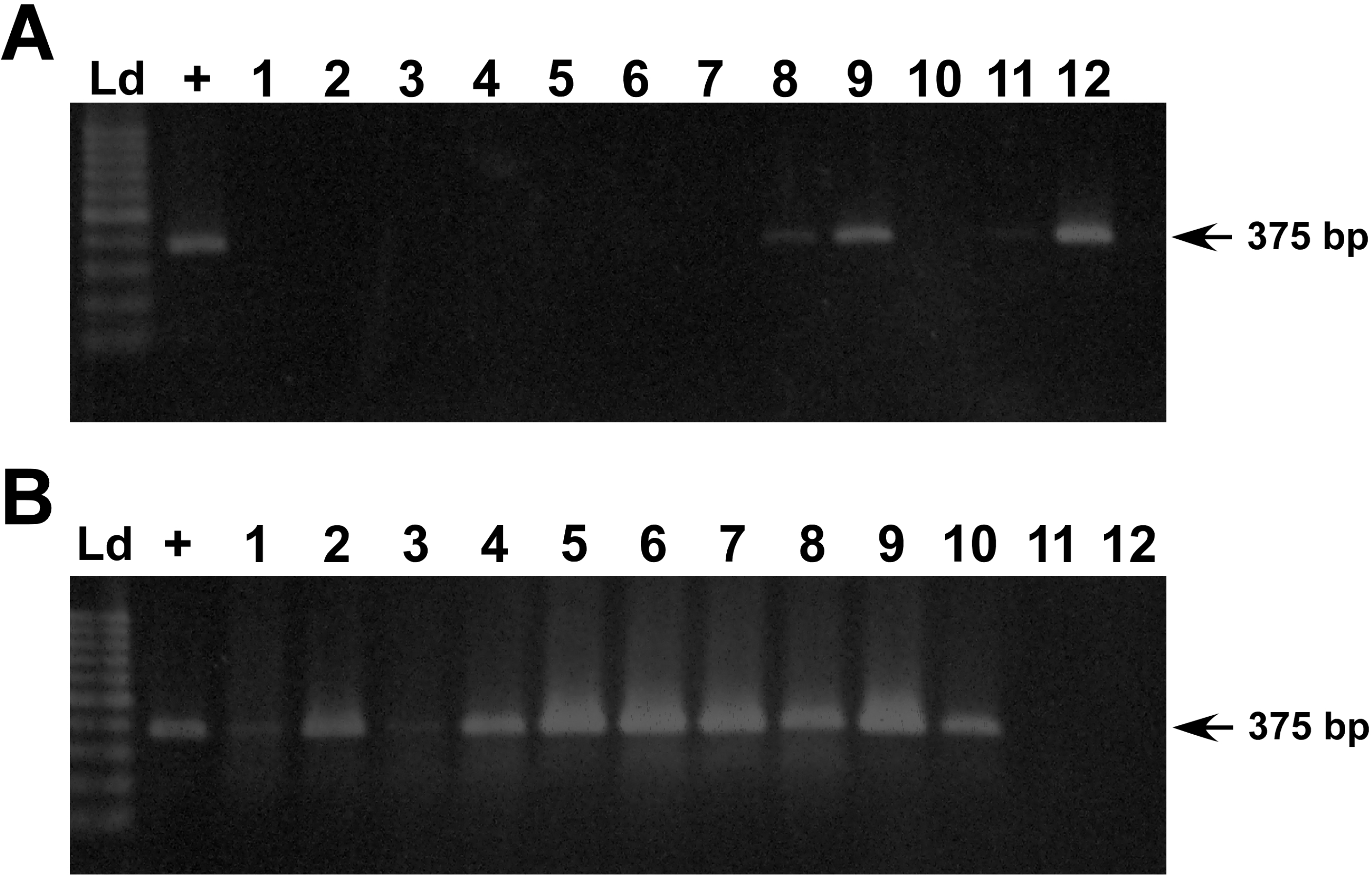
PCR-based detection of *Helicobacter* spp in feces and/or intestine of mice. Randomly selected representative samples of DNA extracted from mice feces (A) or intestines (B). **(A)** Labels: (+) *Helicobacter spp.* DNA positive control; 1-12 = DNA extracted from mice feces; **(B)** Labels: (+) *Helicobacter spp.* DNA positive control; 1-12 = DNA extracted from mice intestine; 1% agarose gel.

**Table 1:** Prevalence of *Helicobacter spp.* infected mice as determined by *Helicobacter* genus-specific PCR analysis.

| Animal<br>mouse | Mouse<br>strain | N°. of mice positive / total number of mice tested<br>(% of positive PCR) |  |
| --- | --- | --- | --- |
|  |  | Feces | Intestine |
| **A** | C57BL/6 | 13 / 22<br>(59%) | 15 / 25<br>(60%) |
| **B** | C57BL/6 | 3 / 5<br>(60%) | 9 / 10<br>(90%) |
| **C** | BALB/c | 11 / 13<br>(84%) | 11 / 13<br>(84.6%) |
| **D** | C57BL/6 | 1 / 7<br>(14%) | 5 / 9<br>(55.6%) |
| **Total** |  | 28 / 47 | 40 / 57 |

## Discussion

The use of living animal models in research requires periodic sanitary screening of animals due to the possibility of undesired infection, which may compromise animal welfare and scientific research. For this reason, laboratory mice should be periodically monitored for parasite, bacteria, fungi and virus, using different methods such as culture, microscopy, PCR and serology (Mähler et al., 2014; FELASA, 2014).

PCR is accepted as one of the most reliable methods for *Helicobacter spp.* detection (Chichlowski and Hale 2009; Casagrande et al, 2010), since culturing these microorganisms requires dedicated infrastructure, laborious and fastidious procedures and also because the presence of several *Helicobacter* species colonizing mice intestine may interfere with bacterial identification (Shames et al. 1995). In addition, serological tests for *Helicobacter spp.* detection present low specificity and may generate false positive results (Whary et al. 2000; Whary and Fox, 2006; Chichlowski and Hale 2009).

Here, we used primers against 16S rRNA able to identify three different *Helicobacter* species that may be associated with several infection processes in susceptible mice strains (Riley et al. 1996). In attention to cross-reactivity issues that could lead to the detection of related microorganisms, first primers sequences were compared to sequence databases from various *Helicobacter* species and then PCR products were sequenced and compared with published sequences of *M. musculus* and *E. coli*. Bioinformatics analysis confirmed that the pair of primers used annealed specifically to conserved sequences of the 16S rRNA gene of *Helicobacter* bacteria.

PCR sensitivity was determined by serial dilutions of DNA extracted from feces and/or intestine. For DNA extraction, we used NaOH or proteinase K instead of commercial extraction kits. Both methods elicited good quality DNA and may work as alternative and less expensive method of DNA extraction for PCR-based screening tests of intestine and feces. PCR product amplification in the expected size was observed using 1 and 10ng of DNA mass. DNA sequence analysis confirmed a similarity of ~95% with the 16S rRNA gene of *H. hepaticus*, *H. bilis* and *H. muridarum*. The screening for bacterial species of the same genus using a single pair of primers is advantageous since it’s minimizes time and costs of the assay (Battles et al. 1995; Shames et al. 1995). It’s important to point out that other *Helicobacter* species have been associated with rodent infection (Hodzic et al. 2001; Fox et al. 2010), therefore depending on the established goals of each animal facility, the choice of primers and PCR strategy for diagnosis may be more challenging. In addition, in some cases the investigation for individual bacterial species identification may be of interest.

Importantly, a high prevalence of *Helicobacter spp.*-infected mice was observed in the facilities screened in this study. Because the prevalence of *Helicobacter spp.* in rodents used for research in Brazil remains largely unknown, the development and application of non-expensive, relatively simple diagnostic tests to routinely screen mice colonies kept in Brazilian research facilities are particularly useful. The successful use of feces is interesting, because it avoids euthanasia, being a useful tool to monitor contamination of small mice colonies. Notably, international councils, such as FELASA and the AALAS, recommend the use of sentinels animals for the health screening routine of rodent colonies (Lipman and Homberger, 2003, FELASA, 2014), while the Brazilian National Council of Animal Experimentation Control (CONCEA) (CONCEA, 2016) does not. Taken into consideration that in Brazil, few companies provide services of microbiological screening tests for laboratory rodents, our findings highlight the need for the development of accessible monitoring routines in order to prevent use of infected animals.

## Acknowledgements

We thank Severino Gomes for technical assistance. The laboratory where this work was performed was supported by grants from the Conselho Nacional de Desenvolvimento Científico e Tecnológico (CNPq), FAPERJ and International Retinal Research Foundation (IRRF) to R.A.P.M.

